# Seasonal challenges of tropical bats (*Rousettus aegyptiacus*) in temperate zones

**DOI:** 10.1101/2021.12.31.473712

**Authors:** Maya Weinberg, Omer Mazar, Lee Harten, Michal Handel, Sophia Goutink, Nora Lifshitz, Gábor Á. Czirják, Yossi Yovel

## Abstract

Egyptian fruit bats (*Rousettus aegyptiacus*) manage to survive and flourish in a large geographic range despite the variability of natural and anthropogenic conditions in this range. To examine the challenges faced by free-ranging *R*.*aegyptiacus* living at the northern edge of their distribution, we performed a retrospective analysis of ∼1500 clinical cases reported by a bat rescue NGO over 25 months, from all over Israel. All cases of injured or stranded bats were evaluated and categorized according to date, place, sex, age, and etiology of the morbidity. The analysis of the data showed an increase in all types of morbidity during the wintertime, with more than twice the number of cases in comparison with the summertime, over two consecutive years. Moreover, we found that the number of abandoned pups peaks during spring till autumn when adult morbidity is minimal. We characterize two prominent types of previously undescribed morbidity in *R*.*aegyptiacus*, one in the form of bacterial illness, and the other associated with feet deformation which affects bats in addition to major anthropogenic-related threats related to synanthropic predators. We analyze the reasons driving winter morbidity and conclude that winter weather and specifically low temperature best explains this morbidity. We hypothesize that *R*.*aegyptiacus*, a fruit-bat of tropical origin is facing major seasonal difficulties near the northern edge of its distribution, probably limiting its further spread northward.

## Introduction

Egyptian fruit bats (*Rousettus aegyptiacus*) are highly common across all Israel which is located close to the northernmost edge of their geographic distribution range (they can be found as north as southern Turkey (1) (see **supplementary Fig 1**). The bats in Israel must thus cope with a relatively cold and wet winter along with many months of hot and dry summer, conditions that are different from the tropical conditions from most of the species’ natural range(2)(3).

It is highly challenging for a tropical species to expand its distribution range deep into the temperate zone and *R*.*aegyptiacus* is the only known fruit bat species to do so(2)(4). This species’ survival demands a year-round availability of fruit, which is enabled in Israel mostly due to anthropogenic plant cultivation. Yet, even with sufficient food, ambient temperature and especially its drop-in winter is a substantial barrier for a tropical species (5) and expanding the geographical boundaries demands copping with that as well. Another major challenge for *R*.*aegyptiacus* in Israel is the very high human population density in Israel (406 inhabitants per km^2^) which results in frequent interactions between bats and humans. As a species that often roost in cities, these bats also suffer from common encounters of synanthropic predators such as cats, crows, and rats (see **Figure 1**A). Despite these challenges, *R*.*aegyptiacus* flourish in urban areas in Israel, mostly due to the widespread availability of deserted structures that are used as roosting places, and thanks to agricultural and ornamental fruit trees which provide a year-round source of fruit (6).

**Figure 1:**
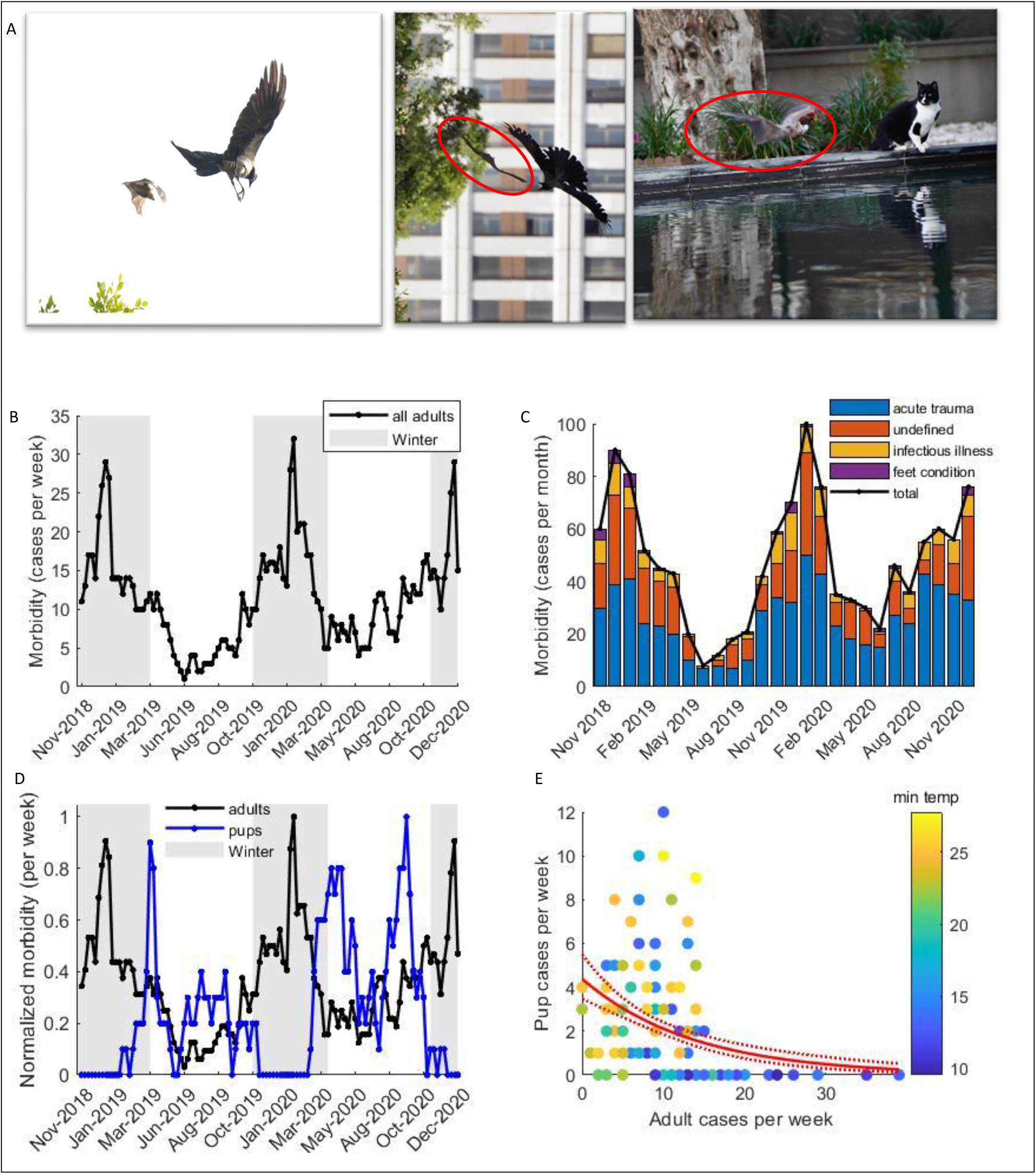
Morbidity in R.aegyptiacus in a temperate area. ***(A)*** *R.aegyptiacus* from urban areas encounter various predators and challenges. *Left and Middle:* A bat threatened by a crow during foraging. *Right*: A bat followed by a stray cat. (Photos in courtesy of Y. Barkai). ***(B)*** The number of adult cases reported during the research period. The graph shows the average number of reported cases per week in two consecutive weeks. Grey shaded areas represent the winter periods. ***(C)*** The proportions of morbidity types in adult cases during the research (acute trauma, illness of infectious origin, feet condition, or undefined). ***(D)*** The normalized number of reported cases per week of adults (black) and pups (blue). ***(E)*** Adult cases as a function of pup cases, the colors present the average minimum temperature per week. The graph depicts a negative correlation between the cases, meaning that as colder it gets more adult cases are reported and fewer to absent cases of pups are reported.

In light of these challenges, we sought to examine the main causes of morbidity of free-ranging Egyptian fruit bats in Israel. It is difficult to obtain reliable morbidity data for wild animals. Most of the information is based on medical reports from rescue centers, wildlife veterinary hospitals, or governmental laboratories (7)(8)(9). Such data is mostly not available for bats. Here, we take advantage of a unique non-governmental organization (NGO) established in Israel in 2016 – ‘*Amutat Atalef*’. This NGO receives and handles hundreds of reports of bats every year. Each bat case is documented including the time and location where it was found, as well as the bat’s estimated age and general health condition based on a first clinical examination. In addition, a picture of the bat is taken *in situ*. By analyzing ∼1500 reported cases arriving over 25 months, we had the opportunity to study the main health issues of *R.aegyptiacus* bats in detail alongside their possible explanatory variables. Our results reveal a clear peak in incidences in winter, which are related to weather conditions but probably also to some food limitations as expected from a bat species living beyond its tropical range.

## Results

We analyzed a total of 1483 morbidity reports received between November 2018 and December 2020. The majority of the calls were for adult bats 1238 (83%). Out of these, 998 (81%) were from urban areas (settlements populated with 30k people and above) and the rest of the calls 240 (19%) were from rural areas, including small villages, nature reserves, and army bases. These differences in numbers most probably reflect the distribution of human reporters. From the 230 cases in which sex was identified, 128 (55%) were females and 102 (45%) males.

There is a significant increase in adult bat morbidity during winter (see **Figure 1**), p<<10^−12^, DF = 109, respectively GLM with a Poisson distribution, explained variable: *number of adult cases per week*, explaining fixed variable: *winter or summer*). The average number of cases per week during the winter was ∼2.1 times higher than during summer (a mean of 15.9 +\- 6 sd with a maximum of 32 in winter, vs. 7.5+\- 3.8 sd, with a maximum of 17 cases only in summer). The winter effect is significant both for rural and urban areas (p<<10^−-12^, p=0.0043, respectively, DF = 109 for both, GLM with a Poisson distribution, explained variable: *the number of cases* in the rural or urban areas, respectively, explaining variable (fixed): winter or summer). We did not find a significant difference between males and females (p=0.071 for sex as the explaining variable, DF =219, GLM with Poisson distribution, explained variable: *number of cases*, explaining fixed variables: *week number, sex*).

An increase in morbidity in winter, compared to summer, was observed in all morbidity types defined. Out of all cases, 69% were categorized while 31% were marked as undefined. Within the diagnosed cases, 79.54% were acute trauma, 17.0% had an infectious origin and 3.5% were attributed as feet chronic, non-infectious condition. 70% of cases with infectious etiology were abscesses with bacterial origin. These proportions among the type of morbidity remained constant year-round (p=0.9, 0.8, 0.72, 0.75, for acute trauma, illness of infectious origin, feet condition, and undefined, respectively, DF =24, GLM models for the explained variable: *rank of cases for morbidity*, explaining variables: *month*, see Methods).

Pup morbidity exhibited different patterns from adults (Figure 1D, blue line). Pup morbidity peaked twice a year in April and September, one month after peak parturition from March and August (10)(11). Pup morbidity was negatively correlated with adult morbidity and temperature: the estimated coefficients - 0.66+\-0.08, -0.25+\-0.04 (mean +\- sd) for GLM model for the *pup morbidity* as the explained variable and the *adult morbidity*, and *averaged minimal temperature* as the fixed explaining variables with interaction, p-values are both > 10^−10^, DF=104, R^2^ = 0.59

Since winter (November-March) in Israel is characterized by relatively harsh weather and by a decrease in food resources (fruit) compared to tropics, in the next step we examined which of these factors better explains the increase in adult winter morbidity. During the Mediterranean winter the precipitation, wind, and temperature are highly correlated (see **Supplementary 2**), therefore we included one weather parameter (the minimal temperature) as a proxy for the weather condition in the models. We found that weather (expressed by minimal temperature) significantly explained morbidity (**Figure 2**A) p<<10^−10^, DF=104, R^2^= 0.57 GLM, with the number of adult cases set as the explained parameter, and *week number* and *minimal temperature* a set as fixed effects, with interaction). Week number and the interaction were also significant with p=2*10^−7^, and p-3*10^−10^, respectively. When fitting a linear curve to the data the influence of temperature on morbidity was estimated as -0.69+/-0.09 cases/deg, meaning that a drop of 5 degrees from 20°C to 15°C increased the number of reports by ∼23 %. Notably, the effect of the temperature on morbidity remained significant even during the wintertime only, proving that the significance is not a result of winter vs. summer batching phenomenon (p= 4.0*10^−5^, DF=44 R^2^= 0.3, GLM with the *number of adult cases* set as an explained parameter, and *week number* and *minimal temperature* a set as fixed effects, *estimated between November first and the end of March only*).

**Figure 2:**
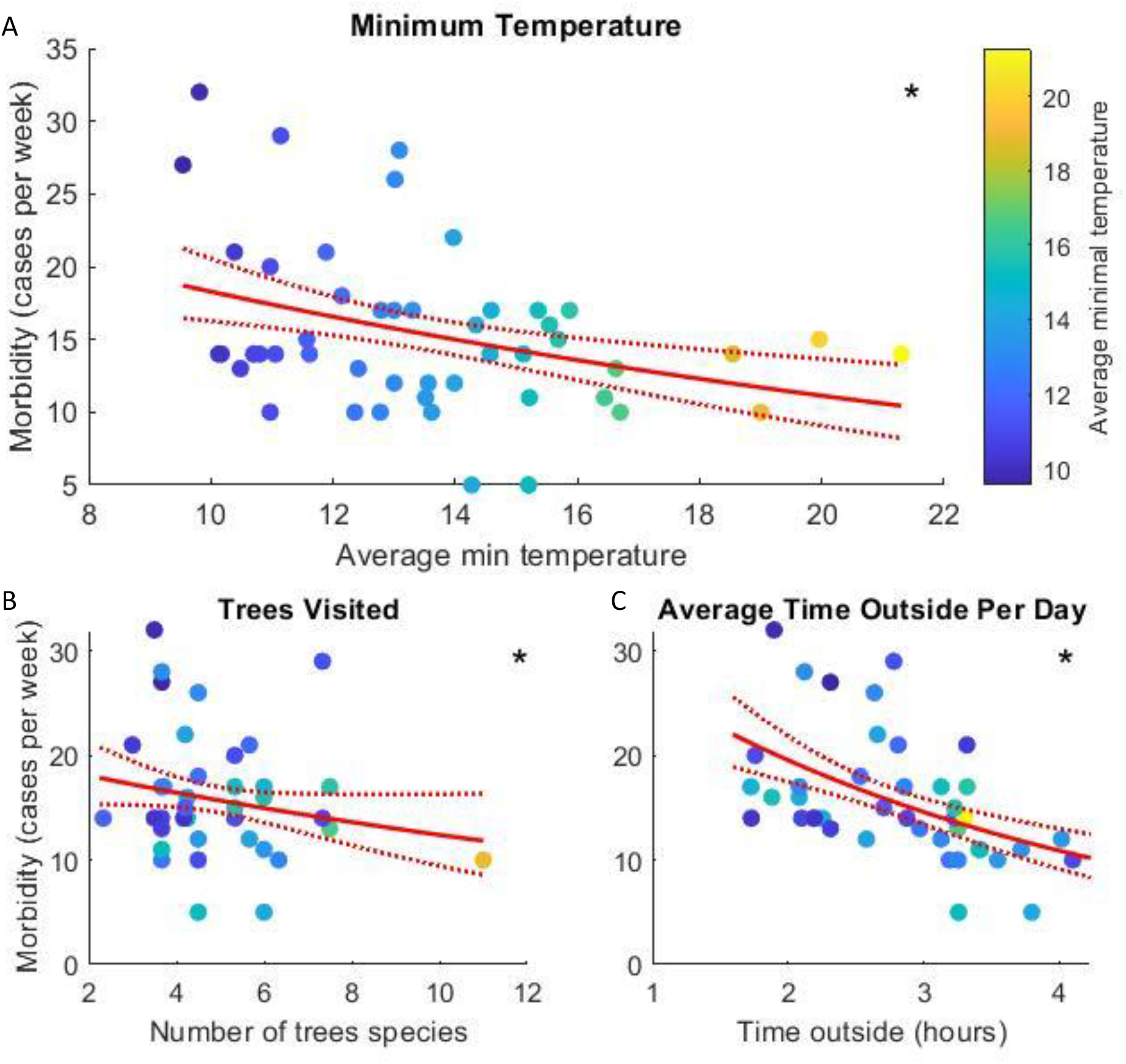
Proximate causes for winter morbidity. Adult cases as a function of (a) minimum daily temperature, (b) number of tree species consumed by the bats, and (c) the average time the bats spent outside their roosting sites. ***In all panels:*** The average minimum daily temperature is depicted by a color code from blue (∼10 degrees) to yellow (∼17 degrees). Each dot represents the mean of two consecutive weeks. Red lines indicate the regression fit and the confidence intervals of the GLM model, with the number of adults’ cases as the explained variable, and each parameter (i.e., temperature, species or time) as the explaining variable, respectively. Since we use a Poisson distribution, the model fits a logarithmic function. Asterisks indicate a significant correlation.

Besides weather conditions, we also tested the effect of fruit availability on adult winter morbidity. As bats visited more fruit trees species during the winter, their morbidity significantly decreased (Figure 2B, p=10^−4^, DF=44 R^2^=0.19, GLM with *morbidit*y set as the explained parameter, and *week number* and *number of fruit species visited* set as fixed effects with interaction). Moreover, as bats spent more time outside the colony their morbidity significantly decreased (**Figure 2**C, p=2*10^−5^, DF=40 R^2^=0.33, GLM, with the number of adult cases set as the explained parameter, and week number and minimal temperature a set as fixed effects, with interaction).

To examine the relative contribution of each factor to morbidity, we analyzed a GLM model with all three predictors (minimal daily temperature, number of tree species, and average time the bat spent outside) set as fixed explaining factors with interactions with the time-in year (week 1-52, for each year). This model explained the results with p=2*10^−7^, DF= 34, and R^2^=0.48. We analyzed which predictor was most important in the model using two methods: Standardized Regression Coefficients, and the Effect on the R-squared when omitting one variable at a time (see statistics in the Methods). According to both of the tests, the temperature is the most informative predictor (see **Table 1**)

**Table 1:**
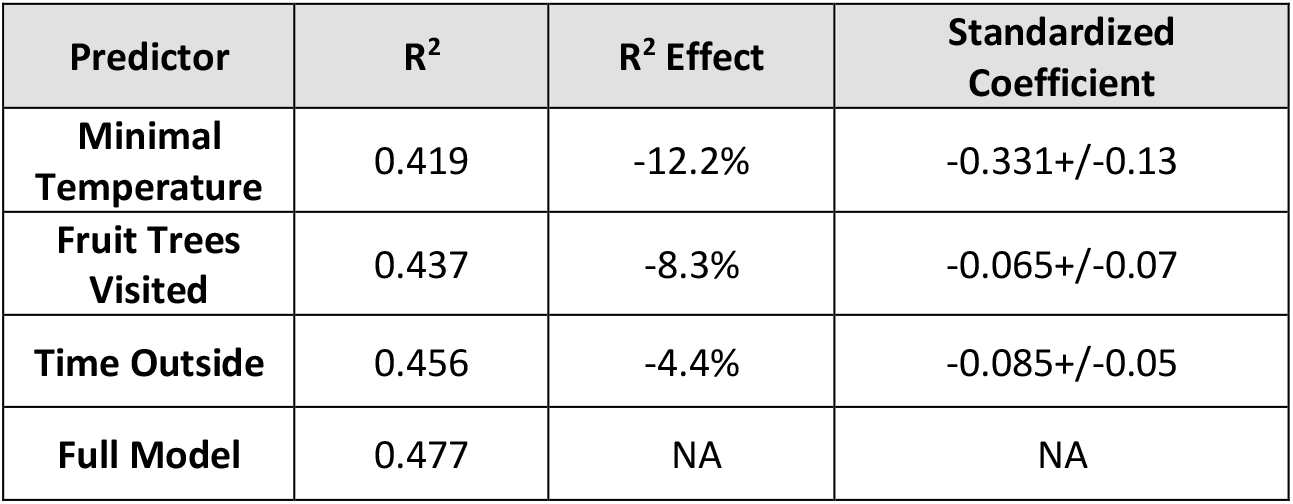
Temperature is the most influential predictor. ***R***^***2***^: the ordinary R-squared when executing a GLM with all parameters except the predictor of each line. ***R***^***2***^ ***Effect:*** the relative effect on R^2^, compared to the full model (bottom line). **All models were significant with p<2*10**^**-7**^ ***Standardized Coefficient***: The coefficient of each predictor (estimation +/-sd) as estimated by a GLM with standardization of all predictors. For both tests, a higher value implies that the predictor has more influence on morbidity than other predictors.

| Predictor | R<sup>2</sup> | R<sup>2</sup> Effect | Standardized Coefficient |
| --- | --- | --- | --- |
| Minimal Temperature | 0.419 | -12.2% | -0.331+/-0.13 |
| Fruit Trees Visited | 0.437 | -8.3% | -0.065+/-0.07 |
| Time Outside | 0.456 | -4.4% | -0.085+/-0.05 |
| Full Model | 0.477 | NA | NA |

We also examined whether the rate of visiting the most visited species (*Ficus microcarpa, Ficus rubiginosa, Melia azedarach, Eucalyptus*, and *Washingtonia*) explained morbidity and found no significant correlation, suggesting that it is not the lack of the main fruit source that drives morbidity (p=0.98, 0.93, 0.97, 0.48, 0.8 respectively, DF=43, n=47, GLM with morbidity as the explained parameter, the minimum temperature and the number of visits at each tree per month as the explaining fixed factor with a Poisson distribution).

## Discussion

Our study aims to elucidate the main drivers and causes of morbidity in free-ranging *R.aegyptiacus* and to reveal the challenges these bats face in temperate climates, beyond their usual tropical habitat (9). Our study is a good example of the value of Citizen Science (12)(13) providing a unique data-set that is extensive and covers a wide temporal and spatial range. Despite the clear benefits, certain limitations exist and these need to be considered (i) our ability to analyze the bat’s precise health status is limited since the data given in the report is often partial. (ii) More bats are found in urban compared to rural areas probably due to differences in human population density. Although over time, human awareness of bats is on the rise, and therefore the number of calls might increase, the yearly numbers of reported bats did not change during the two-year period of this study. Despite these limitations, the data collected by the bat-aid NGO remains the most comprehensive information source regarding *R.aegyptiacus* health in Israel and, to our knowledge, is quite unique world-wide. The relatively high number of cases helps to compensate for these limitations shedding light on the everyday challenges free ranging bats need to cope with.

We found that winter is the hardest time of the year for *R.eagyptiacus*, with a strong correlation between the number of reported morbidity cases and all winter weather parameters we tested (**Supplementary Figure 2**). During the peak of winter, the number of reports for adult bats is twice as high as during the summer, both for the city and rural environments (**Figure 1**B). Notably, this correlation was also observed within the winter months suggesting that it does not result from an overall seasonal difference between summer and winter. A cold winter is known to induce cold stress in a variety of mammals in terms of growth and production (14), reduction in birth weight and enhanced pre-weaning mortality (15), lower immunity abilities (16)(17), and a reduction in basic activity at the cellular level (18). Humans exhibit higher mortality in the wintertime as well (19). Due to long evolutionary history in relatively equable environments, most tropical species and lineages cannot tolerate the abiotic stresses at higher latitudes – especially cold temperature and extreme seasonality – and so they are restricted to the tropics (Brown 2014). They are naturally more susceptible to cold stress symptoms, such as the Florida manatees which suffer from various diseases with prolonged exposure to a temperature below 20°C (20). Several bat species originating from hot climates express torpor upon exposure to cool temperatures (21). *R. aegyptiacus* does not perform torpor (22)(23). Indeed (11) showed that during winter there is a decrease in body mass, followed by an increase during spring and summer, reaching a peak in October, when food availability peaks and weather is still convenient. Therefore it seems that *R.aegyptiacus* are more exposed to all types of morbidity (acute trauma, illness of infectious origin, and feet condition) during the winter season. Living at the edge of their distribution in the neotropics was found to be a stress factor also for the common vampire bat which shows a typical stress leukogram in their white blood cells profile (24). Living on the edge of their distribution is also known to pose difficulties for a variety of other mammals (25). Tropical climates are characterized by monthly average temperatures of 18 °C (64.4 °F) or higher year-round and feature hot temperatures. Novel adaptive traits are required to tolerate stressful abiotic conditions and expand ranges to higher latitudes. The increasing severity of stress acts as a filter, resulting in a decreasing number of species being able to thrive outside the tropics (5).

In our study, adult *R. aegptiacus* were found to suffer from three major types of health issues: acute trauma, illnesses of infectious origin, and chronic feet conditions (**Supplementary Figure 4**). The most common type of morbidity is acute trauma (**Figure 1**C) with almost 80% of reports belonging to this category. Similar results were found for other bats (mostly insectivores)(26)(7)(27)and other wild species, including birds (8)(28). Among these morbidity types, *R.aegyptiacus* mostly suffer from open fractures and wounds related to encounters with urban predators (crows, cats, dogs, rats), and with cars. The second group of morbidity has infectious etiology. Most cases were due to bacterial illness with a typical abscess formation. The abscesses had specific localization (e.g. around the trapezius muscle in the upper back and neck, the soft tissue around the collar bone, and sometimes at the lumbar area) which coincides with the lesion we encountered often in our semi-captive colony (Weinberg et al. unpublished data). Third, the least common cause of morbidity was chronic feet condition. This category included any pathology of chronic origin which affects the tarsal bones and joints. These lesions manifest in an abnormal stretch of the feet that prevents the normal flexion and therefore disables feet usage for hanging. We are acquainted with this pathology in semi-captive bats as well. The last two causes of morbidity have not been previously described in the literature regarding fruit bats, to the best of our knowledge, and demand further investigations. Bacterial illnesses were reported in European insectivorous bats (29)(30). Although *R. aegyptiacus* has been identified as a reservoir for *Bartonella* (31), and several outbreaks of Yersinia pseudotuberculosis have been described in captive colonies (32)(33), none of these were associated with localized abscesses.

Following the immune challenge with bacterial lipopolysaccharide (LPS), sick *R. aegyptiacus* bats “stay home” and avoid foraging trips during the peak of their illness and up to 48 hours later (22). Even as they start foraging post-recovery they do it over a smaller range and during fewer hours. Therefore, it is probable that the number of ill bats found is lower than the actual number in nature, which causes the proportion of injured bats found to be relatively higher. From an infectious biological point of view, bats are mainly examined for viruses (34)(35)(36), although they are highly susceptible to extracellular pathogens such as bacteria (37) and fungi (38) still these pathogens are relatively understudied. Our study provides further evidence, as *R. aegypticaus* develop a profound illness associated with a bacterial origin regularly. We identify this illness as of bacterial origin due to a typical anti-bacterial immune reaction (abscess and pus) as seen in the pictures sent in the group (see an example **Supplementary Figure 4**) and due to previous encounters with this type of morbidity in bats at our semi-captive colony (Weinberg et al. unpublished data). All three identified types of morbidity are present year-round and keep their relative incidence during the year. Undefined cases most probably belong to one of these three categories, but we couldn’t identify them properly due to insufficient information.

The challenges that *R.aegyptiacus* experience in the Israeli winter are also reflected in their parturition seasonality. Parturition completely stops during winter between November to late February (**Figure 1**D), while adult morbidity is at its peak. Only from March onwards did we find lost pups with two clear peaks of incidence in April and September, matching the previously reported pup seasons for this species in the region(11)(10). It seems that *R.aegyptiacus*, in our region divides its two reproduction seasons non-evenly along the year and adopts a different reproduction seasonality that fits the temperate weather conditions, and the fruit availability within its vast range (10)(39). In tropical regions, *R. aegyptiacus* has a single restricted annual breeding season (39) with copulation in July-August, parturition in November-December during the wet summer. Pups are weaned two months later towards the end of the wet season as the climate seems more favorable for foraging. Other fruit bats from tropical regions, with a restricted distribution range, tend to do the same (40).

Although the hardharsg weather conditions in winter seem to be the main factor explaining morbidity, we cannot exclude the role of food limitations. Homoeothermic bats such as *the R.aegyptiacus* need a continuous daily food supply as they don’t have significant fat storage. In insectivorous bats, temporary food scarcities can induce daily torpor even at high ambient temperatures (41)(42). *R.aegyptiacus* is known to be versatile in its diet (43)(6), yet, the consumption of a large variety of fruits during winter, as previously reported, might indicate a shortage in their core staple diet and a need to supplement it with other sources sometimes less nourishing. The reduction in body weight reported during winter (11) can also imply diet limitations. Folivory, for example, was observed mainly during the winter (6).

*R.aegyptiacus* bats tend to stay indoors during bad weather conditions (see **Supplementary Figure 2**). Despite that, we see that the number of reported cases of all morbidity types increases when the weather is unfavorable, which suggests that individuals that do forage outside have a higher chance to be harmed. Previous studies in bats already described the costs of flying in unfavorable weather conditions and show that bats prefer better weather conditions for flight and migration (44). Possible explanations for foraging in bad weather could be that *R.aegyptiacus* must consume food almost on a nightly basis (unlike other bat species which can use daily torpor or hibernate). It is also possible that weaker and hungrier individuals must take the risk and forage under bad weather, and are thus also more prone to morbidity.

*R.aegyptiacus* faces additional challenges in highly populated countries such as Israel. We found that more cases of morbidity are reported in urban areas (79% of cases reports), but this result should be treated with caution due to the reporting bias mentioned above. There is no good estimate of the differences in fruit bat density in urban regions. As a species that thrives in urban environments, they encounter synanthropic predators including humans (and their cars) (45). We observed numerous cases of cat and dog bites and crow attack injuries with 53 cat bite cases described during the examined period and 30 cases of crow attacks. Cat bites are known to injure other bat species as well (26,46) which might lead to systemic bacterial infection with *Pasteurella* sp. In anthropogenic surroundings, cats pose such a prominent threat over bats that behavioral changes of avoidance were examined for the great horseshoe bat, mostly during the reproduction season (47). We have also encountered 15 cases of dog bites and 3 cases of car hits. These findings fit the cause of injuries of other bats in literature(5)(46). The number of fruit bats in urban places in the region is expected to increase as *R.aegyptiacus* is attracted to cities due to the accessible and versatile diet, higher temperature, and man-made roosting structures (48)(49)(50). This phenomenon is probably happening in neighboring countries as Egypt, Jordan, Lebanon, and Turkey, where we know *R.aegyptiacus* roost as well and it might even be happening in the entire range of the species (2). *R.aegyptiacus* is a mammal that thrives in human proximity, yet direct contact with humans is seldom. They keep a clear distance as much as possible during their sleeping time (in caves and deserted buildings, never in rooftops of buildings that are in use) and during foraging time (consuming only ripe fruit and leaves from trees, and never from the ground or market stalls). Even while drinking from manmade pools they go unnoticed (see **Figure 1**A). In this manner, *R.aegyptiacus* represents a very delicate yet successful example of human-wildlife coexistence.

Finally, our study also contributes to the understanding of the factors that determine the distribution range of a species. We demonstrate the challenges faced by a tropical species expanding to a temperate zone. Positioned very close to the northern-most edge of the species’ distribution, *R.aegyptiacus* in Israel has to cope with nearly the most extreme winter in their distribution range (bats in Turkey and Lebanon might suffer slightly colder winters, and indeed lower numbers of *R. aegypticus* are observed there). This harsh winter (relative to their tropical origin) probably limits the distribution range of this species.

## Methods

The fruit-bat NGO ‘*Amutat Atalef*’ received ∼750 reports on fruit bats every year between 2018 and 2020. These messages are conveyed through the “WhatsApp” application, to a group of volunteers who rush to assist the bats. Between November 2018 and December 2020, every text message sent through this group was integrated into a table, which was organized as follows: date, time, location, rural vs urban surrounding, sex, estimated age, clinical description of the morbidity, and/or mortality. All these entries were compared to the provided images and were evaluated blindly and independently by two examiners (MW is a veterinarian doctor specializing in bats for over a decade and SG, a student who was trained by MW) who are well acquainted with *R.aegyptiacus*. A total of 1483 cases have been recorded and analyzed between 4/11/2018 and 20/12/2020. Two cases reported without the location were excluded from the analysis. There is currently no accurate way to determine bats’ age visually, so bats were divided into two age categories: adults or young pups, which were always less than 4 months old, based on visual parameters as described in (11)(51).

Morbidity was categorized into 4 major types of health issues: *acute trauma, illness of infectious origin, feet condition*, and *undefined* (**Supplementary Figure 4**). Morbidity refers to adult bats only, since pups, when found, were all categorized as lost, e.g. involuntarily separated from their mothers. Note, that at this age pups are still dependent on their mothers for supplementary food, thermoregulation, and protection during the day (52). Acute trauma included any type of injuries such as wing fractures, cat bites, crow peaks, car hits, and all other types of recent wounds. *Illness of infectious origin* included all situations of a clear ongoing inflammatory and infectious process such as swollen joints and manifestations of abscesses. *Feet condition* refers to a chronic health situation resulting from an abnormal extension of the tarsal and metatarsal bones that prevents the bats from hanging, eventually leading to exhaustion and death. *Undefined* refers to all cases without sufficient information to allow identification as one of the main categories, or to other conditions. Cases of mortality were reported, in the same manner through the WhatsApp group. Animals were treated in the function of their illness, though some of them died later without notification in the group.

We have collected climate data from the archive of the national meteorological database service (https://ims.data.gov.il/) for the relevant period. Information regarding the daily minimum and maximal temperature and precipitation was collected from the database. Wind speed was collected by the hour and averaged per day. For weekly analysis, the daily data was averaged over the relevant period. Since most cases reported (79%) originate from the urban surroundings in central Israel, we analyzed only the data of the Tel Aviv meteorological station as it represents well urban surrounding climate with its sub climate influence (see urban climate influence over fruit bats.

To track fruit availability and consumption by the bats during wintertime, as well as the amount of time spent outside the roost for commuting and foraging, 12 bats were tracked continuously using GPS from winter to springtime (December 2017 – May 2018) in the Tel-Aviv area (53). The fruit trees visited by the bats including their location and species identification were verified by visiting the actual locations where the bats foraged.

## Statistical analysis

All statistical differences were assessed at a 95% confidence level. As the division for seven days a week is artificial, the analysis was executed after averaging the weekly number of reports with the consecutive week (i.e, using a sliding window with the size of two weeks). Morbidity was defined as the number of cases reported each week or month. It is a countable variable, therefore, we used the Generalized Linear Model (GLM) with the Poisson distribution. The differences between the cases in winter and summer were tested by GLM with *morbidity* as the explained variable and *winter-time* as the fixed explaining variable. Winter was defined as the period between 1\11 and 31\03. Then, we repeated the analysis after dividing the data into rural and urban reports. All GLMs were executed with the Poisson distribution for the number of cases.

The effect of sex on morbidity was analyzed using the GLM, in which the number of weekly reported cases was set as the explained variable and the week-number (i.e. 1-52, for each year) and the reported sex as the explanatory variables.

To test whether the proportions of the causes for morbidity changed in different periods along the year, we clustered the cases into months and ranked them. We then ran a GLM model for the ranked morbidity as a function of the month number in the year (i.e. 1-12, along the research’s period that lasted 26 months).

Because our behavioral and morbidity data-sets were from consecutive years, to evaluate the influence of the time bats spent outside and the variability of the fruit trees the bats visited on morbidity, we duplicated the data collected from the bats in winter 2017-18 to the two consecutive winters of 2018-19, and 2019-20. The trends of the correlations between all pairs of tested parameters during the winter in 2017-18 were similar to the trends during 2019-20 (see **Supplementary 3**). We applied a GLM with the number of cases as the explained variable, and the averaged minimal temperature, average number of trees, while the week-number, the desired variable, and their interaction were set as the explaining fixed variables. To test which of the predictors is the most influential we used two methods: (i) we examined the reduction in R-squared (R^2^) when the variable is omitted from the full model. In this test, we calculated the ordinary R^2^ of the full model then we omitted each predictor and calculated R^2^ again. The R^2^ effect of each predictor was defined as the ratio between the change in R^2^ and the full model’s result. (ii) We compared the predictors’ coefficient in a standardized GLM. First, each predictor was standardized to a z-score according to: 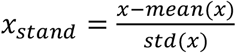. Then, the value of the coefficient was estimated by a GLM model (on all predictors together). The predictor with the largest absolute value for the standardized coefficient is the most influential. All analyses were performed using MATLAB^©^ R2020b.

## Supplementary

**Supplementary Figure 1:**
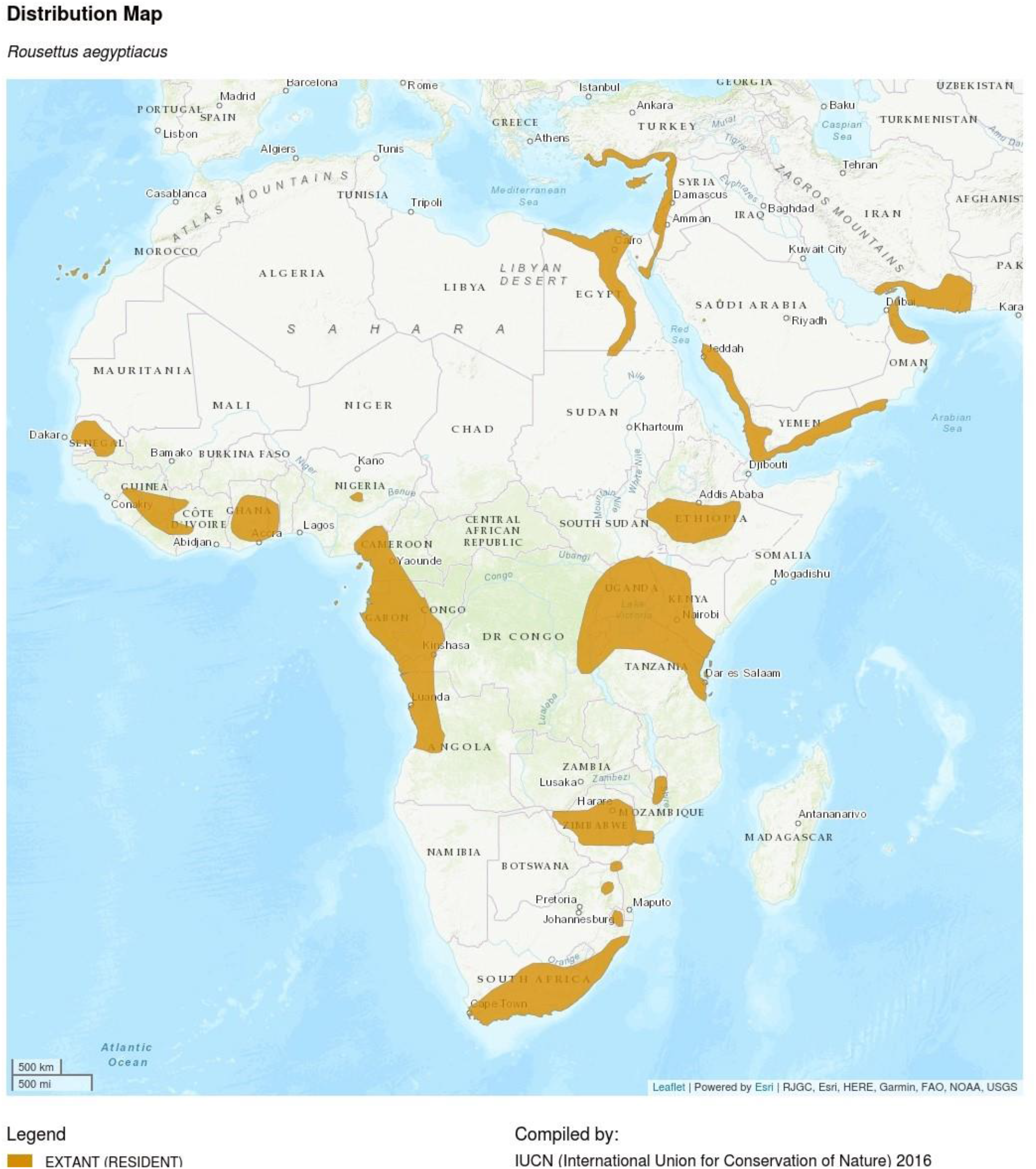
Distribution map of R.aegyptiacus. Origin: IUCN (2016)

**Supplementary Figure 2:**
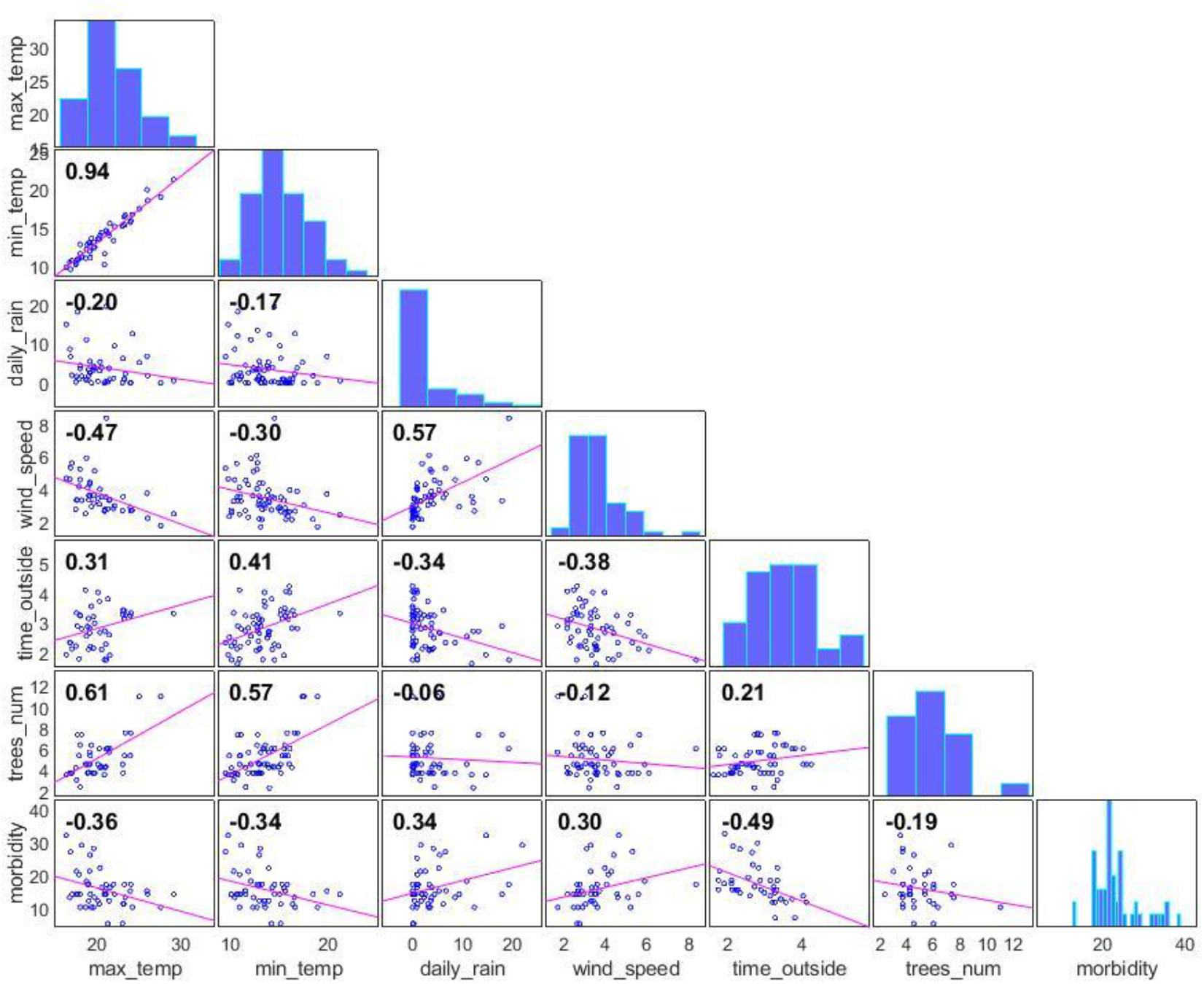
Correlation Matrix between All Parameters during winter (2018-2020). The figure depicts the cross-correlation between each pair of the main measured parameters (averages of the daily data per week, of the following: maximal and minimal temperature, daily precipitation, wind speed, total time of bat outside the cave, number of trees visited, and number of bats cases reported. See methods section for more details). Data regarding the availability of fruiting trees and the time spent outside the cave were collected only during the wintertime, and therefore we present here correlation of this data during the wintertime only. The correlation matrix indicates the following: 1. a strong correlation between all-weather parameters; temperature, wind speed, and precipitation. Therefore, we chose the minimal temperature as the representative parameter for all meteorological data with the highest correlation to the number of adult cases. 2. Weak correlation between the variety of fruiting trees availability and all the weather conditions, indicating the abundance of fruit during the winter. 3. Strong negative correlation between total time spent outside the cave and the weather conditions, implying that as the weather gets worse bats tend to stay indoors. Nevertheless when the weather improves they forage for a longer time (see the correlation between foraging time and minimal temp\ wind speed\ precipitation).

**Supplementary Figure 3:**
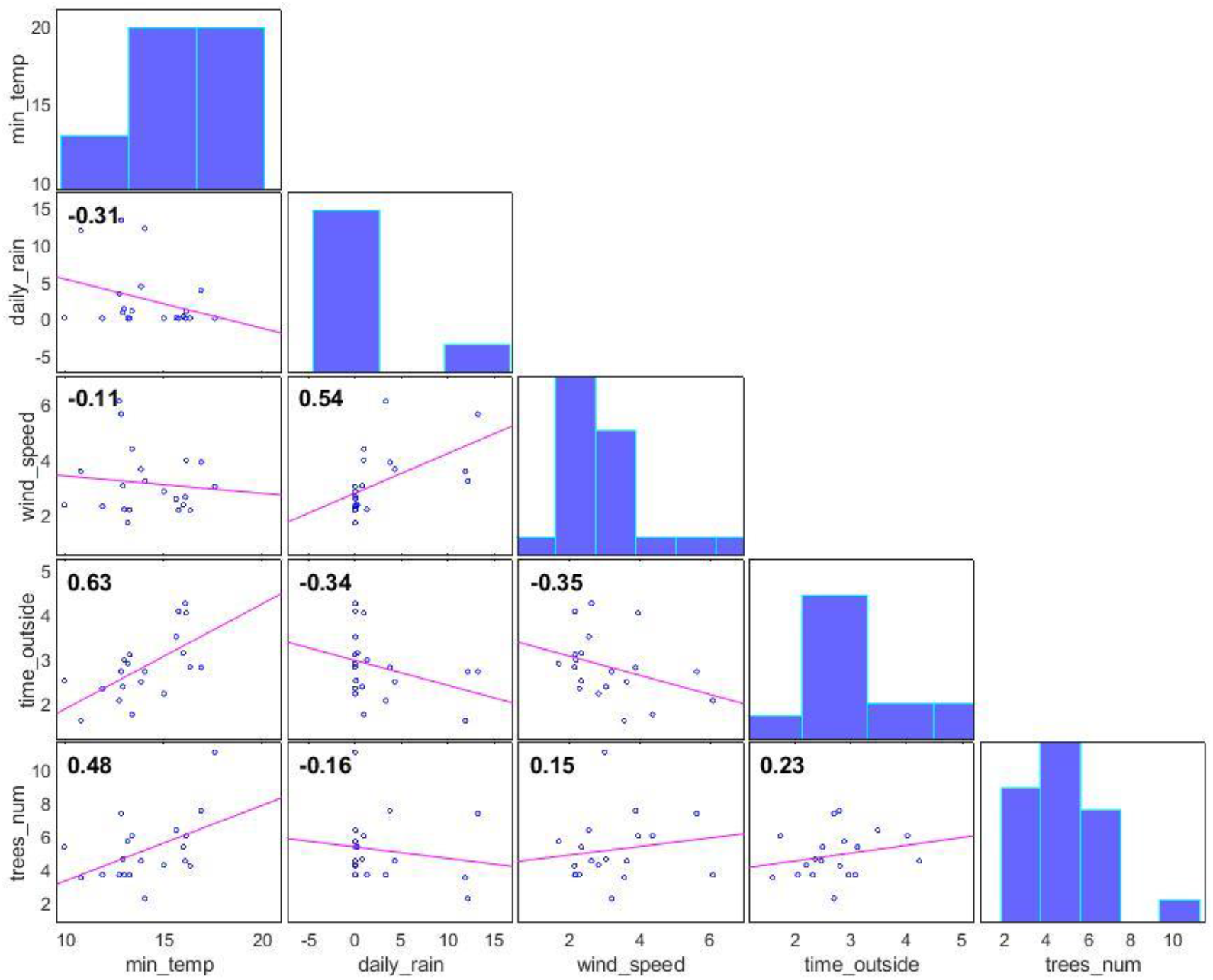
Correlation Matrix between All Parameters during the winter of 2017-2018 only. The figure depicts the cross-correlation between pairs of the data collected during the winter of 2017-1018. The trends of the correlations are similar between the duplicated data and the measured data (the average difference between the trends is 0.037. the standard deviation is 0.14).

**Supplementary Figure 4:**
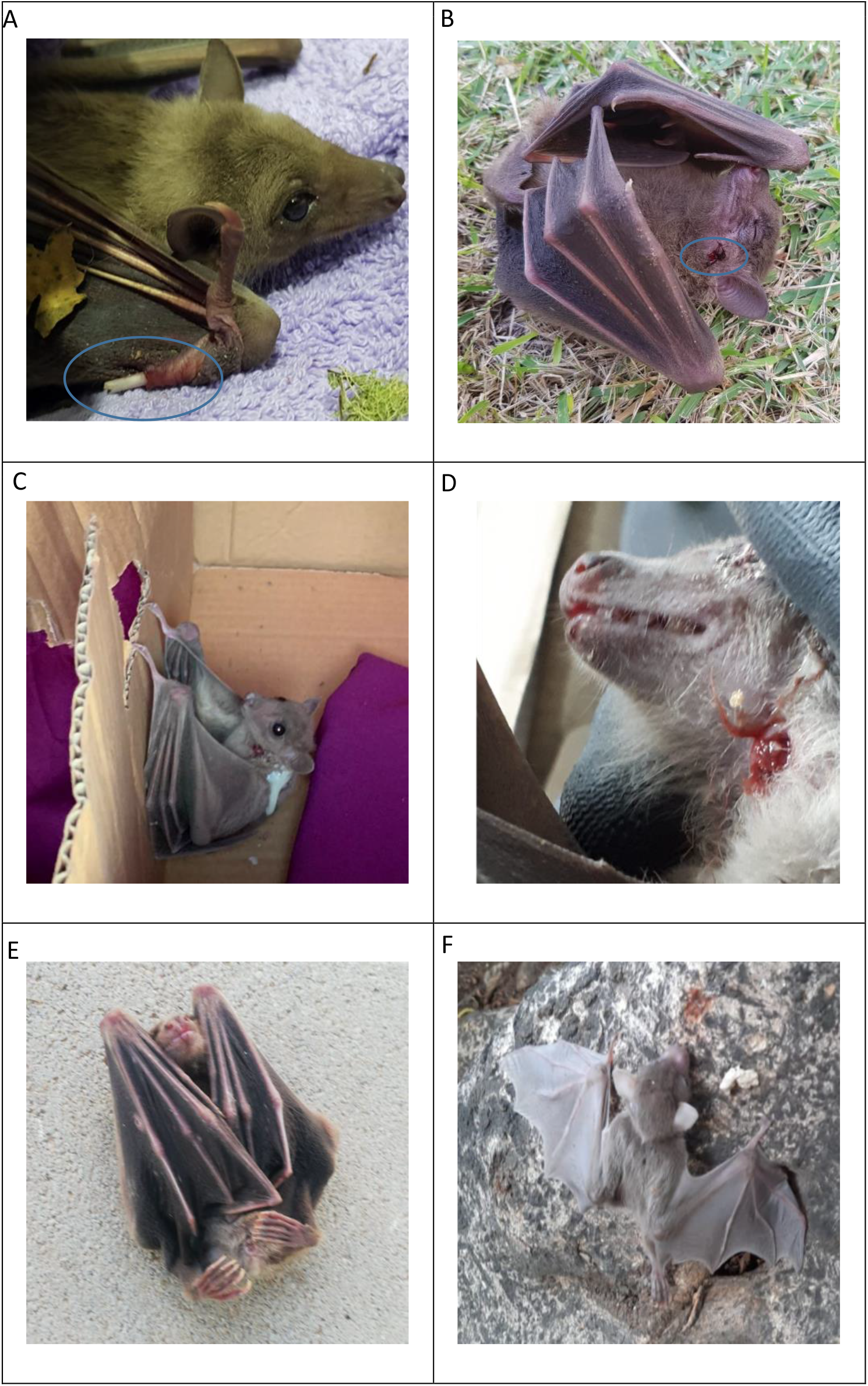
Different types of morbidities as identified from pictures. (A) An open fracture is classified as acute trauma. (B)Pecking by a craw is classified as acute trauma. (C) Pus from a cervical abscess is classified as an infectious illness. (D) Cervical abscess classified as infectious illness (E) Feet condition (F) lost pup. All pictures in courtesy of the Israeli bat sanctuary.

